# Floral microbes provisioned by *Osmia lignaria* establish in larval food stores, but do not affect bee development or survival

**DOI:** 10.1101/2025.08.15.670587

**Authors:** Alexia N. Martin, Clara Stuligross, Neal M. Williams, Helen M. Noroian, Rachel L. Vannette

**Affiliations:** Department of Entomology and Nematology, University of California, Davis, USA; Department of Biology, Kalamazoo College, USA

**Keywords:** Microbial Ecology, Bee Microbiome, Lactobacillus, Floral Transmission, Dispersal, Community Assembly

## Abstract

Microbial dispersal and subsequent establishment among linked habitats can be used to examine drivers of community assembly and function. Flowers host microbial communities that can be acquired and vectored by bees to new flowers, establish within the adult bee gut, and enter food stores (e.g., pollen provisions) of developing larvae. Yet, whether microbes vectored by insects or applied for biocontrol can establish across these habitats and if they affect bee fitness remain unknown. Here, we applied microbes to flowers visited by blue orchard bees (*Osmia lignaria*) and compared microbial communities in flowers, adult bee guts, and pollen provisions before and after inoculation to determine microbial establishment, environmental filtering, and overlap across habitat types. We also inoculated provisions with microbes to test their effects on larval survival and development. Experimentally inoculated microbes were detected in all habitats, demonstrating that flowers are a source of microbial acquisition for adult and larval bees. Additionally, larval health was not impacted by microbe supplementation, indicating tolerance of bee larvae to floral microbes in *Osmia*.

## Introduction

Determining the transmission and establishment of microbial species between habitats is key to understanding their life history and adaptations, and can inform the broader ecological effects of microbial biocontrol application. Like macroorganisms, microbial establishment and subsequent growth depend on dispersal, diversification, selection, and drift (Vellend 2010; Nemergut *et al*. 2013). After dispersal to a new habitat, microbes encounter biotic factors (e.g., priority effects, competition, symbiosis) and abiotic factors (e.g., pH, oxygen, available nutrients) that can act as environmental filters (sometimes also called habitat filters), selecting for species that can survive and proliferate within specific conditions (Kraft *et al*. 2015; Mittelbach and Schemske 2015). However, teasing apart the role of these factors in shaping community assembly and function when using microbial surveys is challenging due to sampling limitations, detection biases, incomplete species knowledge, and variation in dispersal routes (Peay 2014; Kraemer and Boynton 2017). Previous studies have inferred microbial sources using source tracking techniques and algorithms (e.g. Simpson, Domingo and Reasoner 2002; Scott *et al*. 2002; Liu *et al*. 2018; Shenhav *et al*. 2019) or by sampling microbes in potential sources (e.g. Tiusanen, Becker-Scarpitta and Wirta 2024). Even so, directly tracking microbes among habitats is often experimentally intractable in ecologically realistic systems, which prevents us from fully understanding microbial niche breadth and community dynamics. The experimental addition of microbes to a simple system followed by monitoring transmission and establishment across linked habitats can uniquely inform questions of microbial ecology, as well as realistic colonization dynamics and effects on multiple hosts. Here, we use flowers, bees, and pollen provisions (food stores for developing bee offspring) as a tractable model system to investigate the establishment of microbes across uniquely demanding habitats with the same microbial food resources (pollen and nectar).

Flowers host microbial communities within pollen and nectar (reviewed by Vannette 2020). When bees visit flowers to collect these resources for themselves or their offspring, they also acquire microbial communities established in these resources (Herrera *et al*. 2010; Corby-Harris, Maes and Anderson 2014; Rothman *et al*. 2019; Russell *et al*. 2019). Although social bee species exchange microbes through intraspecific interactions (Kwong and Moran 2016), most bee species (∼77%) are solitary (i.e. each female provisions her own offspring) and lack social interactions. As a result, most bees are thought to acquire a majority of their microbial associates from environmental sources that they interact with, like flowers and materials used for nest-building (e.g. soil, leaves, wood, etc.) (Keller, Grimmer and Steffan-Dewenter 2013; Voulgari-Kokota *et al*. 2019; Nguyen and Rehan 2023). It is not entirely understood which environmentally acquired microbes establish in bee-associated habitats (flowers, guts, pollen stores) or how they interact with bees. However, current research suggests environmental microbes may modify floral resources (Herrera, García and Pérez 2008; Vannette and Fukami 2016, 2018; Christensen, Munkres and Vannette 2021) and impact bee nutrition (Dharampal, Hetherington and Steffan 2020), survival (Nguyen and Rehan 2025), development (Dharampal, Danforth and Steffan 2022), and behavior (Good *et al*. 2014; Rering *et al*. 2018; Schaeffer *et al*. 2019; Pozo *et al*. 2020) (reviewed by Martin, Schaeffer and Fukami 2022).

Of the solitary bees that have been studied to date, many show evidence of environmental acquisition of microbes within adult bee guts and pollen provisions. For instance, the gut microbiomes of adult blue orchard bees (*Osmia lignaria*) vary across locations (Cohen, McFrederick and Philpott 2020) and are closely correlated with pollen composition (Vannette *et al*. 2025), suggesting that they are primarily environmentally derived. Similarly, other species of megachilid bees show microbial overlap between flowers and bee guts (McFrederick *et al*. 2017). Microbes identified in the pollen provisions of *O. lignaria* and *O*. *ribifloris* are commonly found in flowers and soil (Rothman *et al*. 2019). Together, these studies strongly suggest that flowers are an important microbial source for both bee gut and pollen provision microbiomes. Yet, the extent to which these microbes successfully disperse between, and establish within, each bee-associated habitat is poorly understood.

Once microbes enter flowers, bee guts, or pollen provisions, they encounter the novel conditions associated with each of these habitats. As a result, microbial species that consistently colonize these habitats can be highly specialized, harboring adaptations to survive, utilize pollen nutrients, and interact with other microbes (Herrera, García and Pérez 2008; Dhami, Hartwig and Fukami 2016; Pozo and Jacquemyn 2019; Alvarez-Perez *et al*. 2021; Christensen, Munkres and Vannette 2021). For instance, a microbe that is adapted to low oxygen environments may be able to establish well in the microaerophilic conditions of a bee gut, but unable to compete in aerobic pollen provisions. However, no studies have directly tracked the movement and subsequent establishment of these specialized microbes into adult bee guts and their pollen provisions.

Here, we leverage a tractable flower-bee-microbe model system to directly trace the establishment and dynamics of a microbial community in multiple bee-associated habitats, as well as the subsequent impacts of these communities on larval health. Specifically, we (1) assessed whether flowers act as sources for microbes in adult bees and pollen provisions, (2) evaluated the role that environmental filtering plays in microbial community establishment across habitat types, (3) examined the extent to which microbial communities overlap between habitat types, and (4) investigated the effect of microbial supplementation on offspring health. To do this, we sprayed an inoculum of bee- and flower-associated microbes onto flowers and then sampled the microbial communities within the flowers, adult *Osmia lignaria* guts, and pollen provisions before and after the treatment. We also applied microbial inoculum directly to pollen provisions and monitored larval survival and development. We hypothesized that flowers would serve as a microbial source for bees and their pollen provisions, but microbial species would vary in their ability to establish among habitat types depending on their initial isolation source (i.e. flower, bee gut, pollen provision). Additionally, if flowers are the only source of microbial acquisition, we expected to find very few microbes unique to guts and provisions. Finally, if microbes within the inoculum positively impact larval bee health, we expected to observe increased survival, increased emergence rates, and less weight loss during development.

## Methods

### Study system

*Osmia lignaria* (Say, 1837), the blue orchard bee, is a solitary, cavity nesting mason bee species native to North America (Figure 1a, Rust 1974). As solitary bees, each female creates her own nest. After locating a suitable nest-site, an *O. lignaria* female collects pollen and nectar from flowers to create a pollen provision, lays an egg on top of the provision, and seals the brood-cell with a mud wall (Figure 1b, Torchio 1989). Each offspring spends its entire larval and pupal stages in this nest, ecloses as an adult in fall and overwinters in diapause within its cocoon. New adults emerge in spring to mate, and females repeat the nesting cycle annually (Torchio 1989). Populations of *O. lignaria* are both wild and commercially managed for use in crop pollination (Torchio 1991).

**Figure 1:**
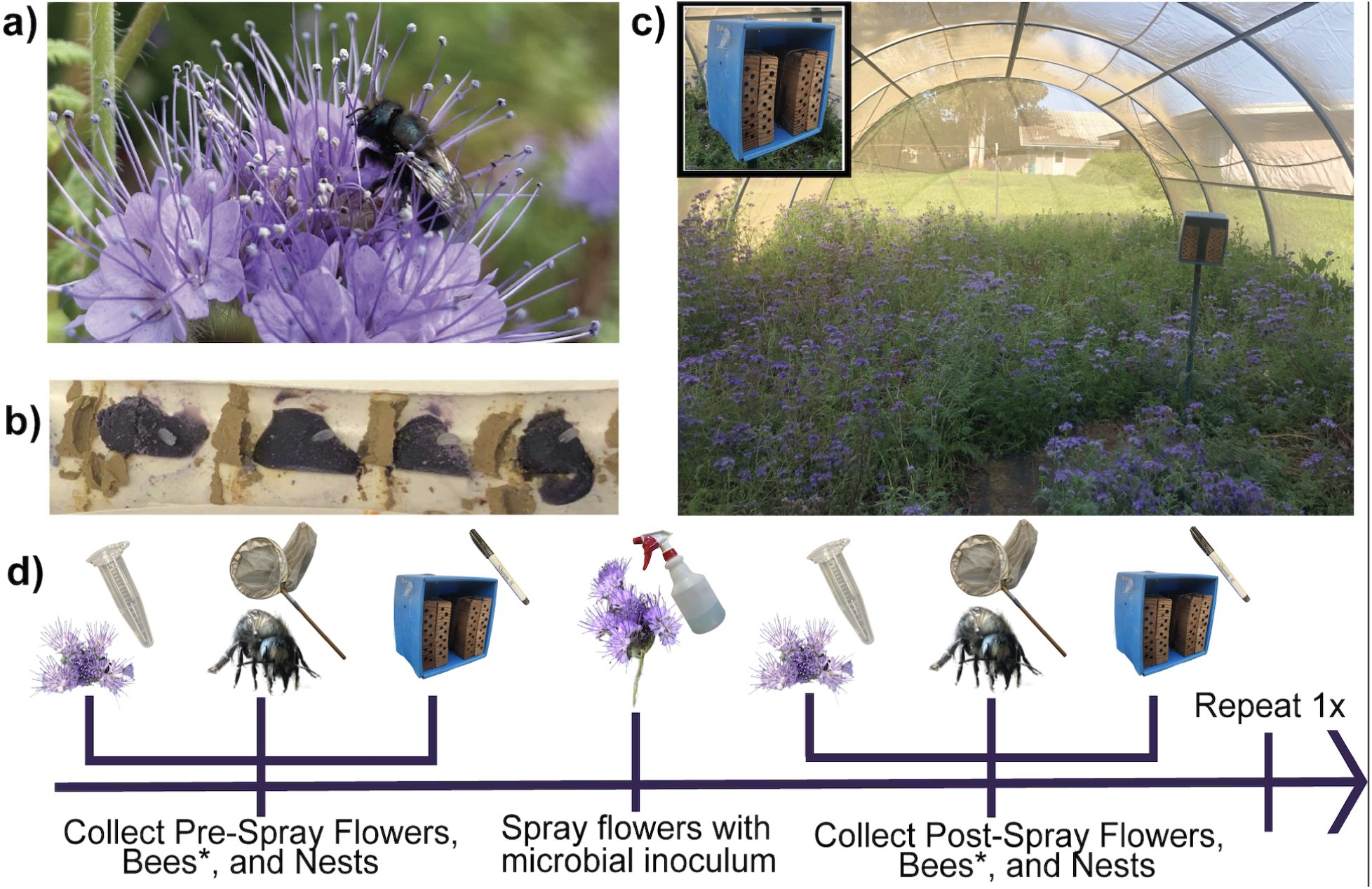
(a) Adult *Osmia lignaria* female on a *Phacelia tanacetifolia* flower (b) Cross-section of a *O. lignaria* nest filled with pollen provisions and eggs, separated by mud walls; (c) Hoop house planted with *P*. *tanacetifolia*. (d) Schematic of sample collection, where all steps were repeated for two rounds except bee collection(*). Photos taken by Alexia Martin and Rachel Vannette.

We performed two experiments: one field experiment to investigate acquisition and community assembly of environmental microbes (Aim 1-3) and one lab experiment to test larval performance in response to microbe addition to pollen provisions (Aim 4). To investigate acquisition and community assembly, six microbes were chosen as focal species to inoculate into flowers (“focal microbes”): two bacteria (*Acinetobacter pollinis*, *Apilactobacillus micheneri*) and four fungi (*Metschnikowia reukaufii*, *Starmerella bombicola*, *Debaryomyces hansenii*, *Aureobasidium pullulans*) (Table 1). These microbes are all frequently found in association with both bees and flowers (McFrederick *et al*. 2017; Alvarez-Perez *et al*. 2021; Rutkowski, Weston and Vannette 2023); however, their primary habitats within these systems remain unclear, and movement between these habitats is poorly understood. Additionally, these microbes were not previously detected in *Phacelia tanacetifolia* flowers under semi-controlled conditions experienced within the experimental hoop houses used for this study (RL Vannette and NM Williams, *Unpublished Data*).

**Table 1:** Focal microbes used to inoculate flowers, their isolation source, the inoculation round they were used, and their recovery in post-spray samples.

| Microbe Type | Microbe Name | Main Isolation Source | Spray Round Used | Recovered in samples? |
| --- | --- | --- | --- | --- |
| Bacteria | *Acinetobacter pollinis* | Flowers | 1, 2 | Yes |
|  | *Apilactobacillus micheneri* | Bee Gut | 1, 2 | Yes |
| Fungi | *Metschnikowia reukaufii* | Flowers | 1, 2 | Yes - through sanger sequencing |
|  | *Debaryomyces hansenii* | Provisions | 2 | Yes |
|  | *Starmerella bombicola* | Provisions | 2 | No |
|  | *Aureobasidium pullulans* | Flowers | 2 | No |
|  | *Vishniacozyma victoriae* | Environmental | 1,2 | Yes |

We chose a community of three bacteria to inoculate into pollen provisions (*Apilactobacillus kunkeei*, *Acinetobacter pollinis*, and *Pantoea agglomerans*). *Apilactobacillus kunkeei* was added because it is a beneficial probiotic in wild bees and honeybees (Nguyen and Rehan 2025; Usta *et al*. 2025), while *P*. *agglomerans* was added due to its usage as a biocontrol agent for fire blight disease (e.g., Stockwell *et al*. 2002; Kim *et al*. 2012).

### Osmia rearing in cages

This experiment was conducted in the spring of 2022 at the University of California, Davis Harry H. Laidlaw Jr. Honey Bee Research Facility in two 6.3 m x 11 m x 2.04 m screen hoop houses. The cage frames were constructed using a wooden base and metal beams, then screened in with a well-fitting fine mesh fabric (Reaco Brand Cage, Santa Rosa, CA). The California native wildflower *Phacelia tanacetifolia* Benth. was grown from the seed bank, as these flowers had been planted within the cages in previous years (Figure 1c; Stuligross and Williams 2020, 2021; Melone, Stuligross and Williams 2024). *Phacelia tanacetifolia* is used by *O*. *lignaria* and provides high-quality pollen and nectar resources (Williams 2003; Boyle *et al*. 2020). When flowers approached full bloom in April 2022, adult male and female *O*. *lignaria* were released into each hoop house. Approximately eight to fifteen females were active at any given time, which was enough to ensure consistent nesting. Bees were provided with wooden nest blocks lined with new paper straws to mimic preferred nest sites and facilitate sampling. Nesting blocks were housed in corrugated plastic shade boxes called “fiesta boxes” (Figure 1c; Boyle and Pitts-Singer 2019). Bees were also provided with a consistent mud source to use for nest construction.

### Microbial suspension preparation and application

To prepare microbial treatments, microbes were plated from glycerol stocks onto Yeast Media Agar (*M*. *reukaufii*, *S*. *bombicola*, *D*. *hansenii*, *Au*. *pullulans*), Tryptic Soy Agar (*Ac*. *pollinis*) or De Man–Rogosa–Sharpe agar + 2% Fructose (*Ap*. *micheneri*). Plates were incubated for 3-5 days at 28 °C. On the day of floral inoculation, microbes were suspended in 1 L sterile water at approximately 4,000 cells/ul per microbe (determined through cell counts on a hemocytometer). The suspension was mixed by gentle inversion and transferred into two clean spray bottles. Microbes were applied directly onto open flowers (approximately 80% of all open flowers) in each hoop house using the spray bottle, with 500 mL of inoculum applied to plant surfaces per hoop house. Individual flowers are open for three days (Williams 1997), with flowers opening sequentially on an inflorescence. Individual bees can live for up to six weeks and presumably vector microbes among individual flowers. The inoculation was performed in two rounds, with Spray Round 1 (approximately 865 million cells/m^2^) occurring on April 12, 2022 and Spray Round 2 (approximately 1.7 billion cells/m^2^) occurring April 26, 2022 and April 28, 2022. Two rounds of inoculation and sampling allowed us to increase replication and examine if focal microbes were maintained in the system after inoculation (likely via bee vectoring among flowers).

### Flower, adult bee, and provision sample collection

Prior to spray inoculation, pooled flower samples (3-4 individual flowers per sample; n=9) and adult female bees (n=6) were collected from both of the hoop houses (Figure 1d). Additionally, completed cells within each nest were marked with permanent marker and assigned as being collected pre-spray inoculation. Pooled flowers (3-4 individual flowers) were collected 1-3 days following each spray round (n=12), and adult female bees were collected 1-3 days following Spray Round 2 (n=12). Nest progress was monitored every 1-3 days for 22 total days with completed cells marked as being created post-spray inoculation. Following completion, nests (n=8) were destructively sampled to obtain pollen provisions, which were assigned as being created pre- or post-spray inoculation (n_pre_=21; n_post_=19).

### DNA extraction and sequencing

Whole guts were dissected from adult bees, and provisions containing egg-2nd instar larvae were dissected from nesting straws using sterile technique. Flower samples were prepared for DNA extractions by adding the pooled flowers to a 5mL tube, covering with 1-2 mL Phosphate Buffered Saline, vortexing for 30-60 seconds, sonicating at 50% power for 20 seconds (Branson Bransonic Ultrasonic Bath 2800), pipetting off the liquid, centrifuging the liquid for 3 mins at 14k rcf to obtain a pellet, pipetting off 0.5-1.5mL of the liquid, and resuspending the pellet in the remaining 500ul of PBS (protocol based on Ushio *et al*. 2015). DNA was extracted from full guts, approximately 1/3 of each pollen provision, and the processed flower samples using the QIAGEN DNEasy PowerSoil Kit according to kit instructions with two modifications. First, the bead beating step was performed on a Benchmark BeadBlaster 24 and included four 20 second cycles. Second, metal beads were added to gut samples during bead beating to improve tissue maceration. The inocula from the second spray round was included as a positive control, and a kit blank was used as a negative control. DNA extract was sent to the Integrated Microbiome Resources at Dalhousie University in Halifax, NS for sequencing of the 16S rRNA (V5/V6) region using 799F (5’-AACMGGATTAGATACCCKG-3’) and 1115R (5’-AGGGTTGCGCTCGTTG-3’) primers (Chelius and Triplett 2001; Anguita-Maeso *et al*. 2022; Christensen *et al*. 2024) and ITS region using ITS86F (5’-GTGAATCATCGAATCTTTGAA-3’; Turenne *et al*. 1999) and ITS4R (5’-TCCTCCGCTTATTGATATGC-3’; White *et al*. 1990) primers via Illumina MiSeq (2x300 bp PE).

### Sequence processing

Sequence data was processed in R (Version 4.2.2 “Innocent and Trusting”; R Core Team 2022) using the standard dada2 pipeline (Callahan *et al*. 2016) to remove primers, merge forward and reverse sequences, and remove chimeras. For the ITS pipeline, primers were removed via ‘cutadapt’ (Martin 2011), and the filtering criteria “maximum errors allotted” was set to 5 for reverse sequences to allow more to pass through the pipeline. Taxonomy was assigned using the Silva database v138.1 (Quast *et al*. 2013) for the 16S data and the Hybrid Unite Database (format 2; Nilsson *et al*. 2019) for the ITS data. Chloroplast and mitochondrial sequences were removed from the dataset using the Phyloseq package (McMurdie and Holmes 2013). Excluding positive and negative controls, an average of 92% of reads per sample made it past quality filtering for bacteria; however, only an average of 8.8% of reads per sample remained after filtering out chloroplasts, mitochondria, and contaminants with most cuts occurring after the mitochondrial removal step. On average, there were 2,169 bacterial reads per sample following filtering. For fungi, an average of 78% of reads per sample made it through both quality and taxonomic filtering with an average of 25,730 reads per sample following filtering. Detailed results from the bacterial and fungal pipeline can be found in the supplemental information. Two sequenced blanks were used with the ‘Decontam’ package (Davis *et al*. 2018) to identify contaminant sequences based on their prevalence across samples and negative controls, resulting in four contaminants removed from the bacterial dataset and no contaminants identified in the fungal dataset. Sufficient sequencing depth was assessed using sampling curves with the ‘rarecurve’ function in the ‘vegan’ package (Oksanen *et al*. 2024). Bacterial samples with fewer than 200 reads (n=11) and fungal samples with fewer than 500 reads (n=2) were removed from their respective datasets (Figure S1). For bacteria and fungi, unassigned ASVs in the top 30 most abundant sequences were manually assigned using NCBI BLAST (Altschul *et al*. 1990) with a 98% cutoff. Any ASVs with multiple equal percentage matches were assigned to the lowest taxonomic level in common between matches. For the fungal pipeline, 42% of ASVs were unassigned at the Kingdom level. To account for this, any ASVs that were present in higher than 0.01% abundance in the entire dataset were manually assigned using NCBI BLAST at a 98% cutoff. Any BLASTed sequences which were assigned as anything other than fungi were removed from the dataset. Because only one (*D*. *hansenii*) out of the four applied fungi was detected via the Illumina amplicon data, PCR and gel electrophoresis were performed on DNA extracted from all of the samples to assess potential bias in the primer set and SnapGene was used to assess primer binding (See Supplemental Methods).

### qPCR to determine bacterial and fungal abundance

Extracted DNA from all sample types was used to quantify bacterial and fungal rRNA copy number via qPCR. To determine the appropriate concentration to use, six representative samples were diluted 1:10, 1:100, and 1:1000. These dilutions, along with the stock concentration, were run and the dilution with Cq values within optimal range (20-30, closest to 25) was chosen. For bacteria, the 1:100 dilution was used for flowers and the 1:1000 dilution was used for guts and provisions. For fungi, the 1:10 dilution was used for guts and flowers and the 1:1000 dilution was used for provisions. For some fungal samples, the final Cq value was outside the 20-30 range, so they were re-run at an adjusted dilution to try to get them into this range. A bacterial standard curve was generated using a plasmid created from the partial 16S rRNA of *Ap*. *micheneri* (LC318485.1), while a fungal standard curve was generated using a plasmid created from the partial 18S rRNA of *Moniliella oedocephalis* (NG_062174.1).

Samples were randomized across plates using the sample() function in R to help account for plate affects. For each bacterial reaction, 5 uL of SYBR, 3.4 uL of PCR water, 0.3 uL of forward primer 799F (10 uM) and 0.3 uL of the reverse primer 1115R (10 uM) were used. Thermocycler conditions were as follows: 95 °C for 3 minutes, followed by 35 cycles of 95 °C for 30 seconds, 52 °C for 30 seconds, and 72 °C for 1 minute. To quantify fungal abundance, the FungiQuant system was used (Liu *et al*. 2012). For each reaction, 5.0 uL PCR Biosystems qPCRBIO Probe Mix, 0.3 uL FQ-F primer (10 uM), 0.3 uL FQ-R primer (10 uM), 0.03 uL of probe, and 3.37 uL PCR water was used. We used the thermocycler conditions described in Liu *et al*. 2012. Each sample was run in triplicate.

### Rearing Experiment

To investigate the impacts of supplemented floral microbes on larval health, we performed a larval rearing experiment from April 2024 to May 2025. The experiment included two treatments: Control (no additional microbes applied) and Microbe Supplementation (*P. agglomerans*, *Ac*. *pollinis*, and *Ap*. *kunkeei* added). We note that two genera in the inoculum are shared with the floral application study above; *P*. *agglomerans* was not applied to hoop houses but is also a common nectar inhabitant. To prepare the Microbe Supplementation treatment, each microbe was first cultured on its preferred medium (*P. agglomerans* and *Ac*. *pollinis* on Tryptic-Soy Agar, *Ap. kunkeei* on De Man–Rogosa–Sharpe agar + 2% Fructose). Next, a bolus of each microbe was added to 2 mL of 30% Sucrose individually, diluted, and counted on a hemocytometer. Glycerol stocks were created by combining all microbes with sterile 15% glycerol to reach a final concentration of 5000 cells of each microbe per dosage (15,000 total cells applied per bee larva). The control treatment consisted of the same volume of 30% sucrose and 15% glycerol, but without microbe supplementation.

The cage set-up described above was stocked with enough adult male and female *O*. *lignaria* to ensure consistent nesting. Nesting progress was checked daily, and completed nests were brought to the laboratory. Within three days of collection, nests were dissected, and non-feeding developmental stages (egg or first-instar larva) were randomly assigned one of the two treatments and inoculated. For each bee, sex was predicted based on its position within the nest and the size of the pollen provision.

Rearing protocols followed Williams *et al*. 2024. In brief, larvae were maintained at 25°C and 60% humidity, with survival and development monitored daily until cocoon spinning was complete. In July 2024, all cocoons were weighed and transferred to gelatin capsules (pierced for air circulation) for pupation monitoring via x-ray. Starting in November 2024, cocoons were re-weighed to determine weight loss through pupation and summer dormancy, then moved into winter conditions (4 °C, 25% humidity). In April 2025, cocoons were removed from winter conditions, re-weighed to determine overwintering weight loss, and transferred to scintillation vials at room temperature (approximately 25 °C). For three weeks, cocoons were checked daily for adult emergence, and sex was noted (Williams *et al*. 2024). For bees that did not emerge, the cocoon was dissected to determine the sex of the bee. If the bee did not develop to the adult stage following cocoon spinning (n=5), the predicted sex was used in subsequent analysis as there was 79% accuracy in correctly predicting sex.

### Statistical analysis

#### General Sample Description

To compare the alpha diversity of microbes among habitat types (flower, bee gut, pollen provision), we used a linear model with microbial Shannon diversity as a response variable and treatment (pre- or post-spray) and habitat type as predictors, with separate models for bacteria and fungi (‘stats’ package, R Core Team 2022). Then, pairwise tests using Tukey’s HSD were performed between each of the treatment types. To compare if microbial inoculation modified the communities found in flowers, adult bee guts or pollen provisions, we used permutational multivariate analysis of variance (perMANOVA) with Bray-Curtis distance using the adonis2 function (‘vegan’ package; Oksanen *et al*. 2024). Separate models were run for each habitat type, with treatment and spray round, their interaction, and hoophouse identity as independent variables.

#### Aim 1: Are flowers a source of microbes for adult bee guts and pollen provisions?

First, the relative abundance of each focal microbe was determined by dividing the number of reads of each focal microbe by the total number of reads within each sample. Then to determine the effect of inoculation on focal microbe presence and relative abundance, we used generalized linear models and assessed significance using chi-squared and F-tests, respectively. Separate models were run for each microbe detected using the amplicon data set (*Ac*. *pollinis*, *Ap*. *micheneri*, *D*. *hansenii* and *V*. *victoriae)* and each habitat type (flower, adult bee gut, pollen provisions). For flowers and pollen provisions, hoop house and the interaction between treatment and spray round were included as fixed effects. For gut samples, only treatment and spray round were included as fixed effects, since guts were not collected before and after each spray. The models for chi-squared tests, in this and all subsequent analyses, were run with family = binomial. The degree of multicollinearity was assessed using the vif() function in the ‘car’ package (Fox and Weisberg 2019), but since all values were below 2, all predictors were retained in the model.

The generated bacterial and fungal standard curves were used to translate the Cq value to log copy number. In order to account for the percentage of reads identified as mitochondrial, chloroplast, or microbial contamination, we combined the qPCR data with the 16S and ITS sequencing results. First, the calculated log copy number was translated into copy number by taking the exponential. Then, for the bacterial data where we used identical primers, the copy number was multiplied by the percentage of reads remaining in each sample after filtering. Finally, we took the log of the adjusted copy number to use in statistical analysis.

To determine if treatment affected the copy number of DNA in the samples, we performed linear mixed models (LMMs) separated by habitat type (‘lme4’ package, Bates *et al*. 2015). The structure of each model was as follows: log(Adjusted Copy Number) ∼ Treatment + (1|Plate ID). We included plate ID as a random effect in order to account for any plate effects. We did not compare copy numbers across habitat types, as we believe that the amount of each sample used for DNA extractions is not directly comparable. To compare the log abundance of fungi and bacteria in each sample, we divided the fungal adjusted log abundance by the bacterial adjusted log abundance to get a ‘microbial abundance ratio’. Values greater than one suggest that fungi were more abundant in the sample than bacteria, while values less than one suggest bacteria are more abundant than fungi. After processing the data, one flower outlier was removed.

#### Aim 2: Does environmental filtering impact microbial establishment across habitats?

To determine if environmental filtering occurred for each focal microbe among habitat types, chi-square and F-tests were performed on a generalized linear model subsetted to only post-spray samples. In these models, inoculated microbe presence or relative abundance was included as the response variables and habitat type was included as a fixed effect.

#### Aim 3: Do microbial communities overlap between habitat types?

To determine the overlap between wild bacterial and fungal communities found among habitat types, inoculated taxa were removed from the data. Next, shared taxa were determined by subsetting the dataset to each specific habitat type and then creating dataframes containing only the taxa shared between habitat types. A Euler diagram was created to visualize the overlap in communities (‘eulerr’ package; Larsson 2024). Finally, we used DESeq2 (Love, Huber and Anders 2014) to identify genera that were differentially abundant amongst habitat types.

#### Aim 4: Does microbial supplementation impact larval health?

We aimed to investigate whether microbial inoculation affected survival, development time, weight loss during development, pupation success, and adult emergence. To assess survival, we performed a chi-square test on a generalized linear model (GLM), with survival status (dead, alive) as the response variable and treatment (Control, Microbe Supplemented) as the predictor variable. To determine the effect on development time, we calculated the number of days it took each larva to progress from their second instar to cocoon completion. We analyzed total development time using a LMM, with treatment and sex (male, female) as fixed effects, and the rearing plate and nest of origin as random effects on development time. To investigate effects on two key developmental stages, we also analyzed time to complete the fifth instar (from defecation to cocoon initiation) and time to spin their cocoon (from cocoon initiation to cocoon completion) using the same model structure as above. To assess weight loss, we calculated (1) total weight lost between cocoon spinning and adult emergence (Total Weight Lost), (2) weight lost during summer dormancy (July-October) (Summer Dormancy Weight Loss), and (3) weight lost during overwintering (October to April) (Overwintering Weight Loss). We analyzed each using a LMM with the same fixed and random effects. Pupation success (yes/no) was analyzed using a GLM with a chi-squared test, with treatment as the predictor. To examine emergence, we performed two models: a chi-square test on a GLM with emergence success (emerged, did not emerge) as the response, and a LMM with days to emergence as the response variable.

## Results

### Microbial diversity and composition vary among habitats and are altered by experimental addition

Alpha diversity differed within bacterial and fungal communities based on habitat type (Bacteria: F = 6.1, df = 2, p<0.01; Fungi: F = 34.8, df = 2 p <0.001), with the highest alpha diversity in flowers (Bacteria Mean = 4.18; Fungal Mean = 2.44), followed by provisions (Bacteria Mean = 3.80; Fungal Mean = 1.95), and then bee guts (Bacteria Mean = 3.08; Fungal Mean = 1.65) (Figure 2a and 2b). Bacterial community composition was highly variable among samples, with few taxa consistently present across all samples within a habitat type; however, fungal communities were much more consistent. Flowers and bee guts were dominated by *Mycosphaerella* and *Cladosporium*, and larval provisions were dominated by those two taxa as well as *Alternaria* and *Podosphaera* (Figure 2c and 2d, Figure 3).

**Figure 2:**
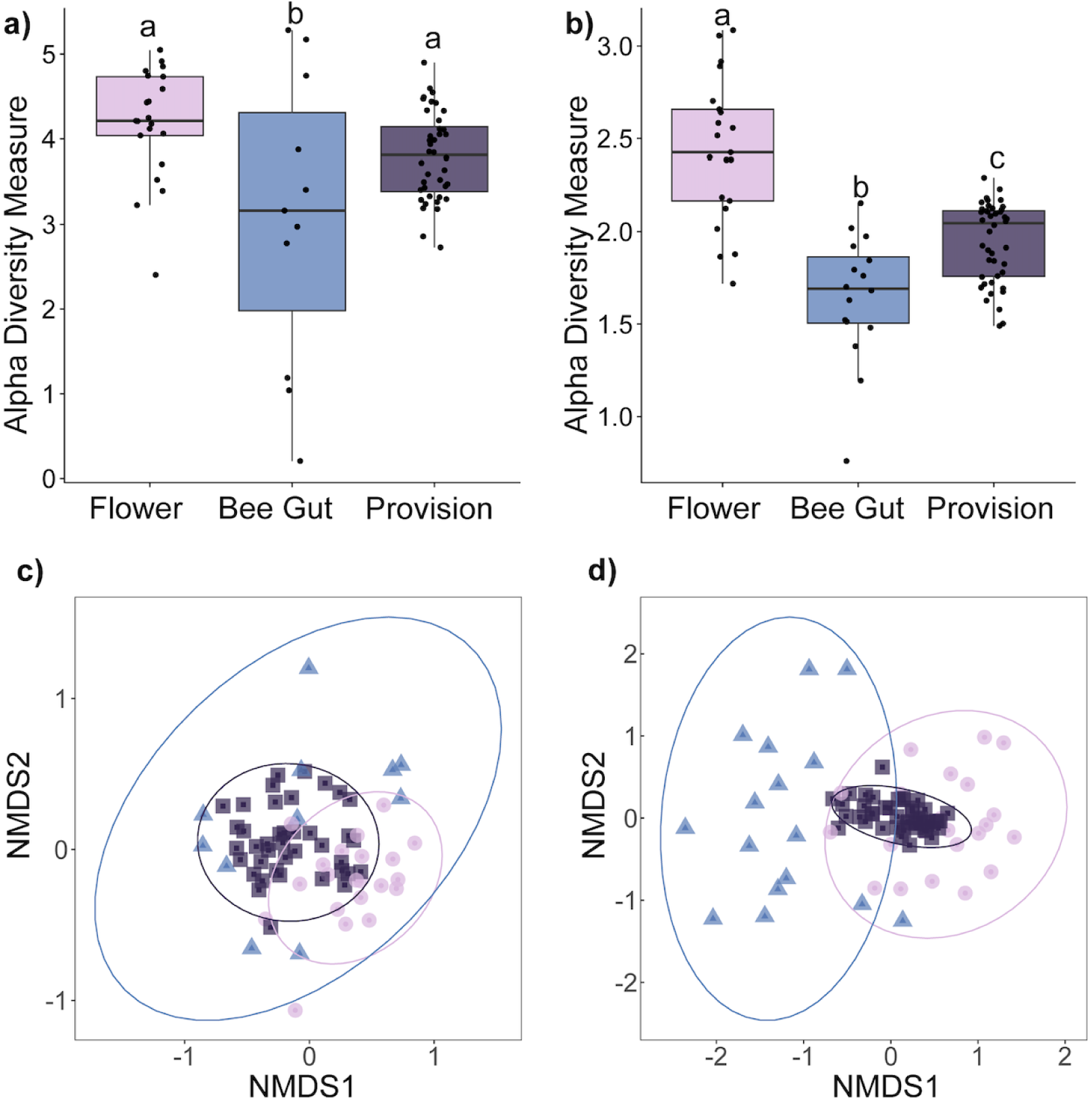
Alpha diversity among habitat types for (a) bacteria and (b) fungi based on amplicon sequencing. Letters signify significant differences between treatments (p < 0.05). PCoA based on habitat type for (c) bacteria and (d) fungi, where light purple circles denote flower samples, blue squares denote bee gut samples, and dark purple squares denote pollen provision samples.

**Figure 3:**
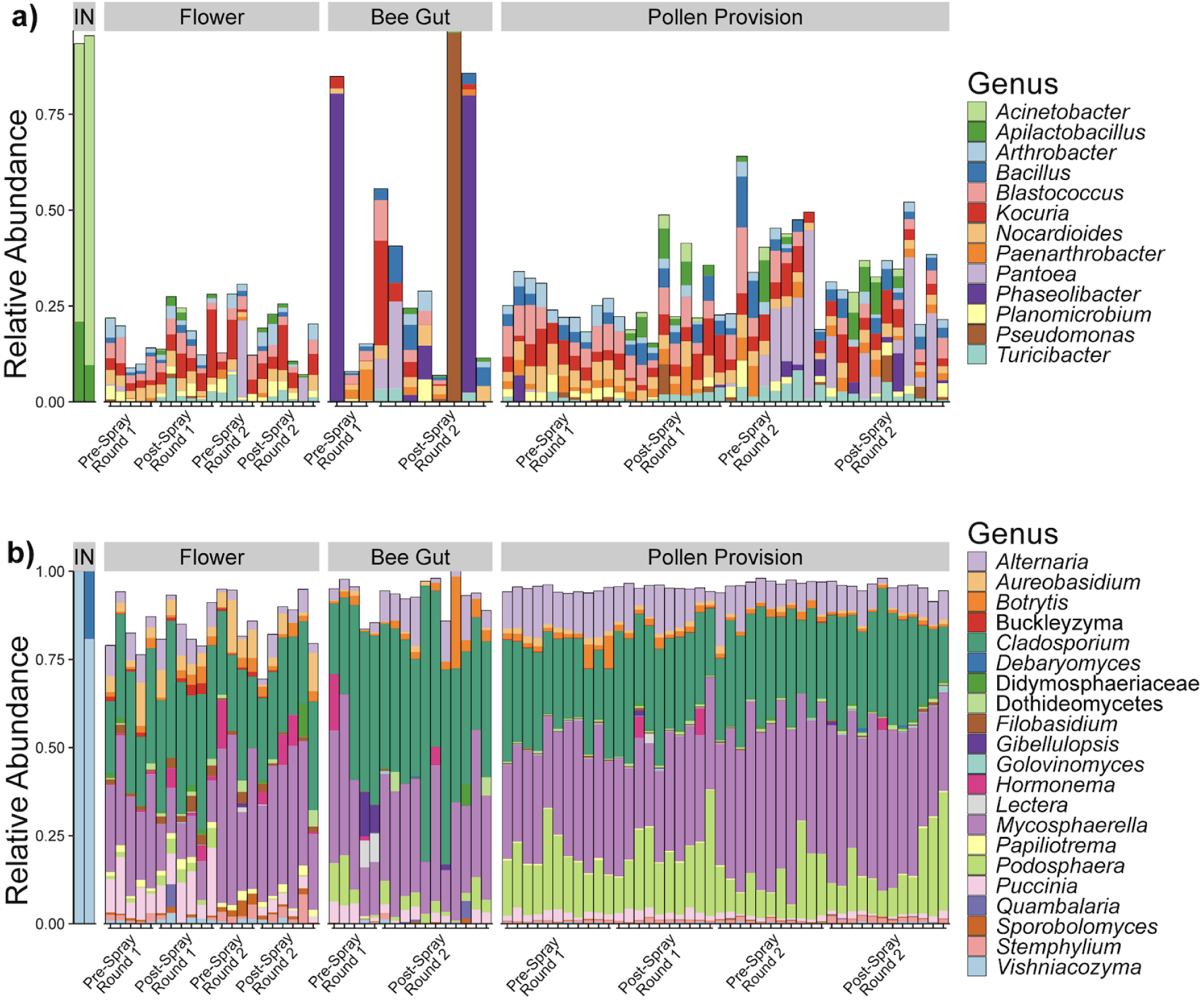
Relative proportional abundance plots of the (a) top 15 most abundant bacterial ASVs and (b) top 30 most abundant fungal ASVs within each sample. Taxa within the plots are at the genus level, though some taxa were included at the class or family level if there was no consensus. “IN” is the inoculum for the second spray round. The white space above bars represents taxa that were found in the sample but were not included in the most abundant taxa.

Experimental inoculation altered microbial communities but only in specific environments, with significant differences detected in fungal communities within flowers and adult bee guts and bacterial communities within pollen provisions (Tables 2 and 3). Bacterial communities in pollen provisions were also influenced by the spray round, cage, and the interaction between spray round and treatment. Fungal communities in flowers and pollen provisions were affected by spray round. Additionally, some groups differed in variance for the fungal communities. This included adult bee guts between treatments (betadisper F = 7.4, df = 1, p = 0.011) and pollen provisions between spray rounds(betadisper F = 5.4, df = 1, p = 0.027).

**Table 2:** Permanova examining effects of treatment, spray round and hoophouse on bacterial community composition within each habitat. Results Permanova table based on a Bray-Curtis Distance Matrix for the 16S (bacterial) data.

| Habitat Type | Effect | DF | Sum of Squares | R2 | F | Pr(>F) |
| --- | --- | --- | --- | --- | --- | --- |
| Flower | Treatment | 1 | 0.42 | 0.059 | 1.23 | 0.094 |
|  | Spray Round | 1 | 0.43 | 0.059 | 1.24 | 0.062 |
|  | Cage | 1 | 0.44 | 0.062 | 1.29 | 0.056 |
|  | Treatment x Spray Round | 1 | 0.37 | 0.052 | 1.087 | 0.286 |
|  | Residual | 16 | 5.51 | 0.77 |  |  |
|  | Total | 20 | 7.18 | 1.00 |  |  |
| Gut | Treatment | 1 | 0.60 | 0.14 | 1.46 | 0.11 |
|  | Cage | 1 | 0.50 | 0.11 | 1.21 | 0.26 |
|  | Residual | 8 | 3.27 | 0.75 |  |  |
|  | Total | 10 | 4.36 | 1.00 |  |  |
| Provision | Treatment | 1 | 0.58 | 0.049 | 2.10 | **0.006** |
|  | Spray Round | 1 | 0.74 | 0.062 | 2.67 | **0.002** |
|  | Cage | 1 | 0.42 | 0.036 | 1.53 | **0.045** |
|  | Treatment x Spray Round | 1 | 0.50 | 0.042 | 1.83 | **0.017** |
|  | Residual | 35 | 9.65 | 0.81 |  |  |
|  | Total | 39 | 11.90 | 1.00 |  |  |

**Table 3:** Permanova examining effects of treatment, spray round and hoophouse on Fungal community composition within each habitat. Results of Permanovas based on a Bray-Curtis Distance Matrix for the ITS (fungal) data.

| Habitat Type | Effect | DF | Sum of Squares | R2 | F | Pr(>F) |
| --- | --- | --- | --- | --- | --- | --- |
| Flower | Treatment | 1 | 0.28 | 0.088 | 2.035 | **0.043** |
|  | Spray Round | 1 | 0.39 | 0.12 | 2.80 | **0.012** |
|  | Cage | 1 | 0.23 | 0.071 | 1.65 | 0.11 |
|  | Treatment x Spray Round | 1 | 0.087 | 0.027 | 0.63 | 0.81 |
|  | Residual | 16 | 2.21 | 0.69 |  |  |
|  | Total | 20 | 3.19 | 1.00 |  |  |
| Gut | Treatment | 1 | 0.58 | 0.24 | 4.40 | **0.007** |
|  | Cage | 1 | 0.14 | 0.055 | 1.014 | 0.46 |
|  | Residual | 13 | 1.74 | 0.71 |  |  |
|  | Total | 15 | 2.47 | 1.00 |  |  |
| Provision | Treatment | 1 | 0.12 | 0.032 | 1.58 | 0.18 |
|  | Spray Round | 1 | 0.54 | 0.14 | 7.00 | **0.002** |
|  | Cage | 1 | 0.089 | 0.023 | 1.14 | 0.27 |
|  | Treatment x Spray Round | 1 | 0.048 | 0.013 | 0.62 | 0.57 |
|  | Residual | 39 | 3.024 | 0.79 |  |  |
|  | Total | 43 | 3.83 | 1.00 |  |  |

### Aim 1: Focal Microbes were detected in flowers, adult bee guts, and pollen provisions suggesting that flowers are a source of microbes

Of the six microbes that were sprayed onto flowers, only three were detected using amplicon sequencing, including both bacteria *Ac*. *pollinis*, *Ap*. *micheneri*, and a single fungal species (*D*. *hansenii)*. Additionally, the environmental fungus *Vishniacozyma victoriae* was overwhelmingly detected in the inoculum despite not being purposefully included. We do not know where *V. victoriae* originated, but because it is present within the environment it may have been present in the spray bottles (Rush *et al*. 2022; Kristjuhan, Kristjuhan and Tamm 2024). Regardless, it was included in the analysis because it was applied to the flowers. The yeasts *M*. *reukaufii*, *Au*. *pullulans*, and *S*. *bombi* were not detected in the inoculum or any of the samples, potentially due to primer bias (See Supplemental Methods). This bias could have also been a reason why *V*. *victoriae* seemed so abundant within our inoculum. Of the four inoculated microbes that were detected, none were present in any samples prior to the first spray (Figure 4).

**Figure 4:**
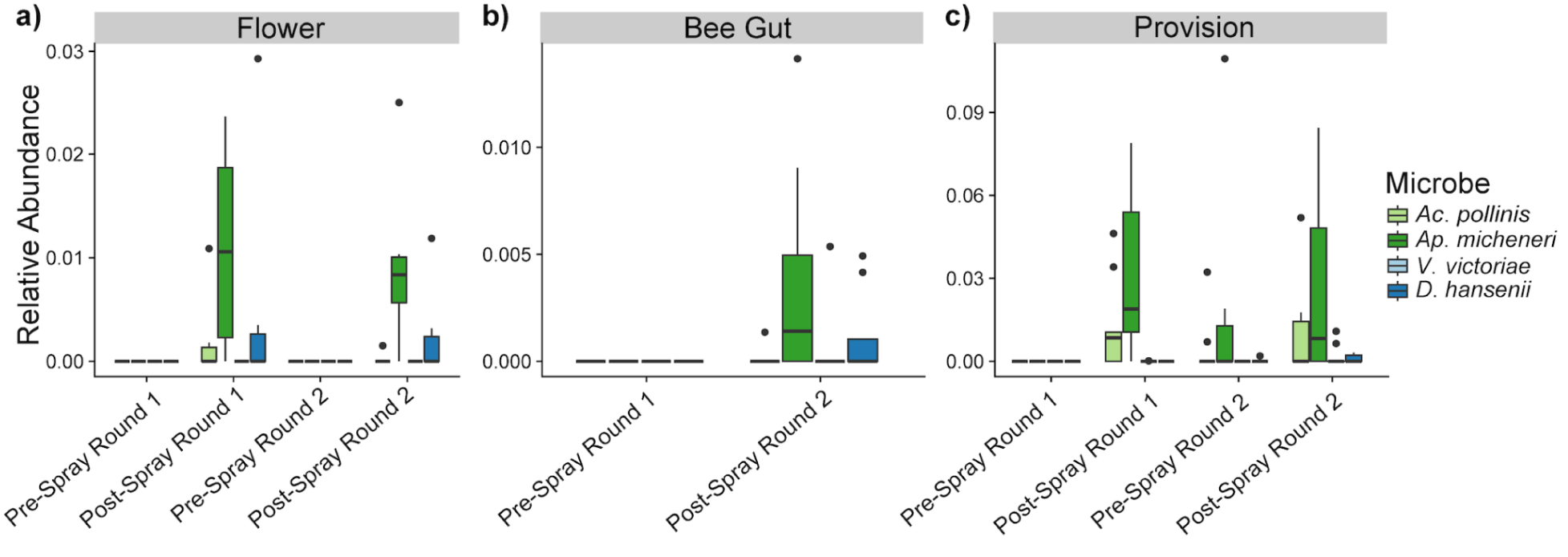
Relative proportional abundances of focal microbes across the three sampled habitat types (flowers, adult bee guts, and pollen provisions). *Acinetobacter pollinis* and *Apilactobacillus micheneri* were used in both spray rounds, while *Debaryomyces hansenii* was only used in spray round 2. *Vishniacozyma victoriae* was found in the fungal inoculum.

The presence and relative abundance of *Ac*. *pollinis* increased in provisions following flower inoculation, with relative abundance 5.38 times higher following inoculum application (Tables 4 and 5). Its establishment differed between spray rounds, increasing in presence after the first round and remaining in flowers before and after the second round. The presence of *Ap*. *micheneri* increased in flowers, adult bee guts, and provisions following inoculation, with establishment in provisions differing between spray rounds (Tables 4 and 5). There was also a 3.83-fold increase in relative abundance of *Ap*. *micheneri* within provisions, while it was only detected in guts and flowers following inoculation. The establishment and relative abundance of *V. victoriae* within pollen provisions only differed based on spray round and cage (Tables 4 and 5); however, there was no effect of treatment. Finally, *D. hansenii* was not found in any flowers pre- or post-spray but was detected in 1/11 post-spray guts. Additionally, *D. hansenii* was detected in only 4.5% of pre-spray provisions (1/22) and 18% of post-spray provisions (4/22); however, this difference was not statistically significant.

**Table 4:**
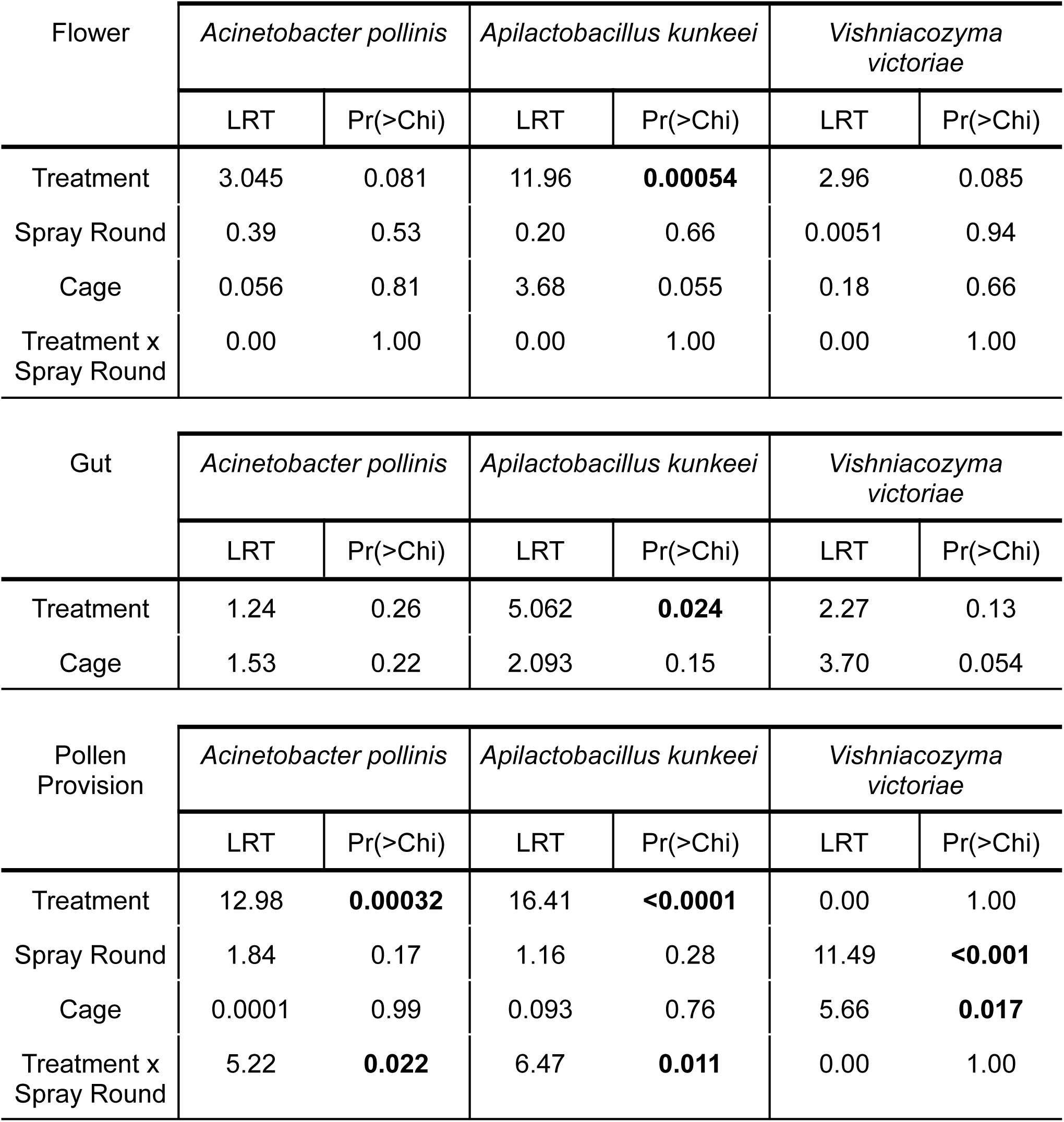
Chi-square tests for examining differences in presence of *Acinetobacter pollinis, Apilactobacillus micheneri, and Vishniacozyma victoriae* across habitat types (flowers, adult bee guts, and pollen provisions).

| Flower | *Acinetobacter pollinis* |  | *Apilactobacillus kunkeei* |  | *Vishniacozyma victoriae* |  |
| --- | --- | --- | --- | --- | --- | --- |
|  | LRT | Pr(>Chi) | LRT | Pr(>Chi) | LRT | Pr(>Chi) |
| Treatment | 3.045 | 0.081 | 11.96 | **0.00054** | 2.96 | 0.085 |
| Spray Round | 0.39 | 0.53 | 0.20 | 0.66 | 0.0051 | 0.94 |
| Cage | 0.056 | 0.81 | 3.68 | 0.055 | 0.18 | 0.66 |
| Treatment x Spray Round | 0.00 | 1.00 | 0.00 | 1.00 | 0.00 | 1.00 |

| Gut | *Acinetobacter pollinis* |  | *Apilactobacillus kunkeei* |  | *Vishniacozyma victoriae* |  |
| --- | --- | --- | --- | --- | --- | --- |
|  | LRT | Pr(>Chi) | LRT | Pr(>Chi) | LRT | Pr(>Chi) |
| Treatment | 1.24 | 0.26 | 5.062 | **0.024** | 2.27 | 0.13 |
| Cage | 1.53 | 0.22 | 2.093 | 0.15 | 3.70 | 0.054 |

| Pollen Provision | *Acinetobacter pollinis* |  | *Apilactobacillus kunkeei* |  | *Vishniacozyma victoriae* |  |
| --- | --- | --- | --- | --- | --- | --- |
|  | LRT | Pr(>Chi) | LRT | Pr(>Chi) | LRT | Pr(>Chi) |
| Treatment | 12.98 | **0.00032** | 16.41 | **<0.0001** | 0.00 | 1.00 |
| Spray Round | 1.84 | 0.17 | 1.16 | 0.28 | 11.49 | **<0.001** |
| Cage | 0.0001 | 0.99 | 0.093 | 0.76 | 5.66 | **0.017** |
| Treatment x Spray Round | 5.22 | **0.022** | 6.47 | **0.011** | 0.00 | 1.00 |

**Table 5:** Results of F-tests for examining the relative abundance of *Acinetobacter pollinis* and *Apilactobacillus micheneri* across the three habitat types (flowers, adult bee guts, and pollen provisions)

| Flower | *Acinetobacter pollinis* |  | *Apilactobacillus kunkeei* |  | *Vishniacozyma victoriae* |  |
| --- | --- | --- | --- | --- | --- | --- |
|  | F-value | Pr(>F) | F-value | Pr(>F) | F-value | Pr(>F) |
| Treatment | 1.65 | 0.22 | 13.23 | 0.0022 | 2.66 | 0.12 |
| Spray Round | 1.50 | 0.24 | 0.049 | 0.49 | 0.93 | 0.35 |
| Cage | 0.40 | 0.53 | 5.077 | 0.039 | 2.33 | 0.15 |
| Treatment x Spray Round | 0.21 | 0.65 | 1.22 | 0.29 | 0.49 | 0.49 |

| Gut | *Acinetobacter pollinis* |  | *Apilactobacillus kunkeei* |  | *Vishniacozyma victoriae* |  |
| --- | --- | --- | --- | --- | --- | --- |
|  | F-value | Pr(>F) | F-value | Pr(>F) | F-value | Pr(>F) |
| Treatment | 1.14 | 0.32 | 1.058 | 0.33 | 1.61 | 0.23 |
| Cage | 1.33 | 0.28 | 0.0085 | 0.93 | 2.92 | 0.11 |

| Pollen Provision | *Acinetobacter pollinis* |  | *Apilactobacillus kunkeei* |  | *Vishniacozyma victoriae* |  |
| --- | --- | --- | --- | --- | --- | --- |
|  | F-value | Pr(>F) | F-value | Pr(>F) | F-value | Pr(>F) |
| Treatment | 5.15 | **0.030** | 5.76 | **0.022** | 0.0051 | 0.94 |
| Spray Round | 0.32 | 0.57 | 0.12 | 0.73 | 13.96 | **<0.001** |
| Cage | 1.85 | 0.18 | 0.82 | 0.37 | 5.096 | **0.030** |
| Treatment x Spray Round | 0.93 | 0.34 | 1.31 | 0.26 | 3.62 | 0.065 |

Despite not being detected in the inoculum or any samples in the Illumina sequencing data, *M. reukaufii* was detected using *Metschnikowia-*specific primers (See Supplemental Methods). It was found in both inocula (using gel electrophoresis only), 7 flower samples (one pre-spray, six post-spray), 6 bee gut samples (none pre-spray, six post-spray), and 14 provision samples (three pre-spray, eleven post-spray) (Supplemental Figure 2). Additionally, the presence of *M*. *reukaufii* significantly increased with treatment for both guts (p<0.05) and provisions (p<0.05). *Metschnikowia reukaufii* presence did not differ across habitat types.

We also found support for our hypothesis that some focal fungi did not appear in the sequencing data due to primer binding issues (See Supplemental Methods). For the inoculated fungi that were present in the sequencing dataset (*V*. *victoriae* and *D. hansenii*), SnapGene (SnapGene software) was able to detect primer binding sites for both the forward and the reverse primers and no differences in the forward primer and microbial sequence. For *V*. *victoriae*, there were six base differences when the reverse primer was aligned and for *D. hansenii* there were three base differences. On the other hand, the focal fungi that did not appear in the sequencing data only had the forward primer bind (*Au. pullulans*) or no primers bind (*S*. *bombicola*, *M*. *reukaufii*). When primers and the microbial sequences were aligned, *Au. pullulans* had no differences in the forward primer and five base differences in the reverse primer, *S*. *bombicola* had two differences in the forward primer and seven differences in the reverse primer, and *M*. *reukaufii* had two base differences for the forward and reverse primers.

We did not detect a significant effect of treatment on microbial abundance measured by the adjusted log copy number for bacteria nor fungi in any of the habitat types (Supplemental Figure 3). However, fungi were more abundant than bacteria in all samples (flower: 2.73, bee gut: 2.46, pollen provision: 2.36).

### Aim 2: Inoculated microbes differed in relative abundance across habitats, suggesting environmental filtering occurs

To investigate if environmental filtering occurred, presence and relative abundance of each inoculated microbe was compared across habitats. Both *Ac*. *pollinis* and *Ap*. *micheneri* differed in relative abundance across habitats (*Ac*. *pollinis*: F_2,37_ = 3.6, p = 0.04; *Ap*. *micheneri*: F_2,37_ = 4.0, p = 0.03), attaining the highest relative abundance in pollen provisions. No differences in presence or relative abundance were detected for *V*. *victoriae* or *D*. *hansenii*.

### Aim 3: Most ASVs were unique to one habitat type, though flowers and provisions shared the greatest proportion of microbes

For the bacteria sequenced and found in at least one sample (2,643 total ASVs), 74% of ASVs were unique to one of the three habitat types (Figure 5a). The highest unique bacterial richness occurred within flowers (43%). Of the 26% of bacterial ASVs shared between habitat types, the highest shared richness (9%) occurred between flowers and provisions and the lowest shared richness occurred between bee guts and flowers (4%). For the fungi sequenced and found in at least one sample (1,483 total ASVs), 79% of

**Figure 5:**
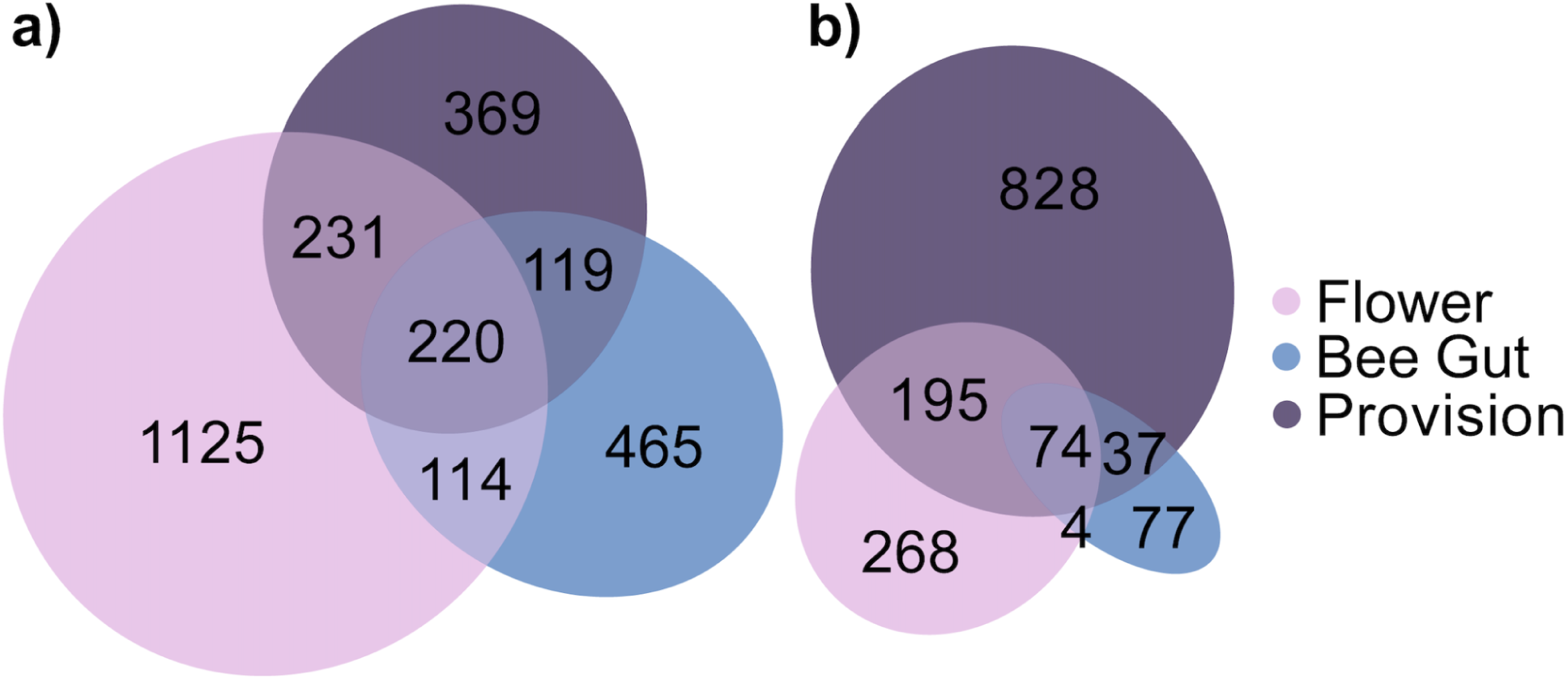
Total (a) bacterial and (b) fungal ASVs unique and shared between flowers, adult bee guts, and pollen provisions.

ASVs were unique to one of the habitat types with the highest unique richness within pollen provisions (56%) (Figure 5b). Similar to bacteria, the highest shared richness of fungal ASVs occurred between flowers and provisions (13%) while flowers and adult bee guts shared the fewest microbes (0.3%). For both bacteria and fungi, multiple genera were found to be differentially abundant (FDR<0.05) across the habitat types though there were more differentially abundant taxa for fungi (n=20) than bacteria (n=4) (Supplemental Figure 4 and 5). The highest differential abundances were found between provisions and guts for bacteria and in provisions for fungi.

### Aim 4: Microbial supplementation had little impact on most bee health metrics

Microbial supplementation of pollen provisions did not significantly affect larval survival (p=0.38). Although developmental timing differed by sex--females took longer to complete the fifth instar (t = -2.6, df = 120.57, p < 0.01) and complete total development (t = -7.4, df = 113.41, p < 0.001)--treatment did not affect any of the analyzed developmental stages (Fifth Instar: t = 0.27, df = 94.28, p = 0.78; Cocoon Spinning: t = 0.74, df = 93.40, p = 0.46, Total Development: t = -1.17, df = 91.22, p = 0.25) (Figure 6a). Similarly, weight loss during development varied by sex (Summer Dormancy: t = 12.8, df = 110.2, p < 0.001; Overwintering: t = 6.4, df = 121.4, p < 0.001; Total Weight Loss: t = 11.27, df = 121.67, p < 0.001) (Figure 6b). Treatment had a small but significant effect on weight loss during summer dormancy (t = 2.01, df = 86.04, p < 0.05), but not during overwintering (t = -0.18, df = 83.2, p = 0.86) or overall (t = 0.95, df = 85.13, p = 0.35). Nearly all larvae successfully pupated (131/135), with no effect of treatment on pupation success (p = 0.95). Finally, microbial treatment did not impact emergence success (χ^2^: p=0.73), but both sex and treatment influenced emergence timing. Males emerged 1.3 days earlier than females, and bees in the microbe-supplemented treatment emerged almost one day later than controls (Sex: t = -3.81, df = 60.26, p < 0.001; Treatment: t = 2.62, df = 2.64, p < 0.05) (Figure 6c).

**Figure 6:**
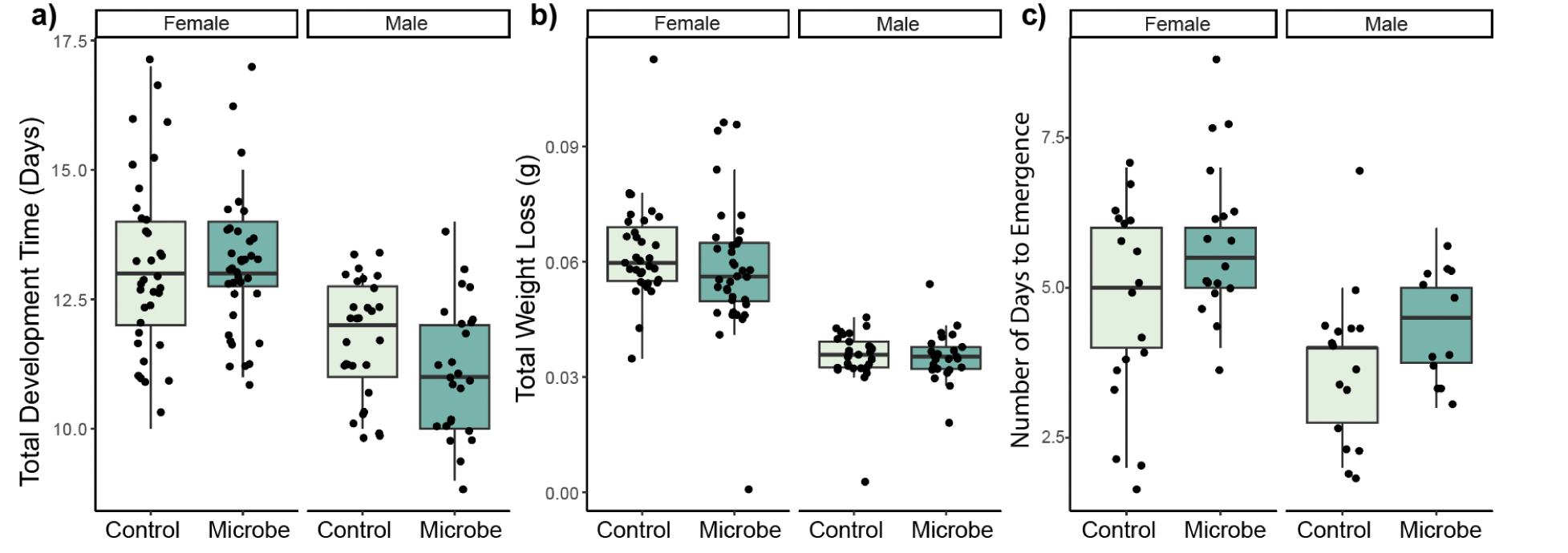
Microbial effects on larval health metrics based on treatment (control, diet microbe supplementation) and confirmed offspring sex (female, male) (a) Total number of days it took for the offspring to develop from first instar to cocoon spinning. (b) Total weight lost between summer dormancy and adult emergence in October 2024. (c) The number of days it took for larvae to emerge after cocoons were warmed to room temperature.

## Discussion

Our controlled field experiment provides clear empirical evidence that flowers are a source of microbial acquisition for bee guts and pollen provisions, but microbial communities are subsequently filtered among pollen provisions and adult bee guts.

Consistent with previous studies on the gut microbiota of *O. lignaria*, we found that the microbial ASVs in adult bee guts and pollen provisions reflect the microbial ASVs present on flowers (Cohen, McFrederick and Philpott 2020). Adult *O*. *lignaria* have very low microbial read counts at adult emergence (Crowley and Schaeffer 2024), which suggests that these microbes are not carried over from their larval stage, during which they consume the pollen provision. Together, these results support the growing body of literature that adult *O*. *lignaria* acquire their microbiome from environmental sources. Yet, our data show that flower microbes differ in their ability to establish across bee-associated habitats.

Unlike social bee species that transfer established microbes between individuals (Kwong and Moran 2016), establishment of microbes in the adult female gut does not seem to be a requirement for establishment within the pollen provision. Here, we saw that the wild bee-associate *Ap*. *micheneri* established within mother bees and pollen provisions, while the flower-associate *Ac*. *pollinis* only established within pollen provisions. This suggests that females may serve as a vector for floral microbes to pollen provisions without needing to harbor them within their gut microbiomes (Mundt and Hammer 1968; Vellend 2010; Duar *et al*. 2017). To create a pollen provision, the female mixes nectar regurgitated from the frontmost portion of the gut (the crop) with pollen groomed from her scopa (a structure on the exterior of the abdomen; Torchio 1989). The crop is frequently filled and emptied of nectar as bees create their pollen provisions, and previous studies on honeybees indicate that it is nearly devoid of established bacteria (Martinson, Moy and Moran 2012). Additionally, the nectar is regurgitated soon after intake, so it likely does not pass to the hindgut where most of the bee gut microbiome is established (Martinson, Moy and Moran 2012). As a result, microbe-rich nectar may be solely transferred into the provision and floral-specialized microbes may not establish in the bee gut. As such, provision microbes are not necessarily able to be predicted by gut-associated microbial communities of free-flying bees. Similarly, pollen carried on the scopa is not ingested by the mother and likely serves as another microbial source for the pollen provision but not the bee gut. Here, we saw that the wild bee-associate *Ap*. *micheneri* established within mother bees and pollen provisions, while the flower-associate *Ac*. *pollinis* only established within pollen provisions.

Although bacterial communities were extremely variable among and within habitats, the fungal communities were remarkably consistent. The presence of the same fungal taxa within and across habitat types could be because these taxa are core to the plant species, because they were dominant in the experimental hoop house environment, or because they are endophytic within the pollen (Hodgson *et al*. 2014). Notably, a recently published geographic survey of *O. lignaria* pollen provision across California found high relative abundances of different fungal taxa (*Mycosphaerella* and *Cladosporium*) in pollen provisions but strong associations between fungal and pollen composition (Vannette *et al*. 2025), suggesting fungal acquisition from pollen. In addition to their consistency, qPCR data revealed that fungi were more abundant than bacteria. Although we did not examine interactions between them, bacteria and fungi co-occur in each habitat, such that they may inhibit, facilitate, or not impact each other’s growth. For instance, the common floral bacterium *Asia astilbes* can decrease growth of the yeast *M*. *reukaufii* (Rering *et al*. 2020). As a result, it is possible that the variation and low abundance of bacterial communities could be due to competitive dominance by the fungi in each habitat. Additional experiments investigating the co-growth of microbes in bee-associated habitats may offer further insight into this quandary.

The general decrease in alpha diversity from flowers to provisions to guts suggests that environmental filtering of microbial communities occurs between these habitat types. Abiotic and biotic factors of each habitat may influence which microbes survive and reproduce in each habitat, such that increased strength of filtering could reduce microbial diversity (Nemergut *et al*. 2013; Mittelbach and Schemske 2015; Moran, Ochman and Hammer 2019). Each of the investigated habitats has unique conditions and can be considered difficult for microbes to colonize in. Floral nectar contains high sugar, low nitrogen, and antimicrobial chemical compounds, which can act as an abiotic filter (Adler 2000; Huang *et al*. 2012). Subsequently, microbes which are able to inhabit nectar may undergo an additional biotic filter through direct and/or indirect interactions with other microbes that occupy the same niche, which may lead to competition or mutualisms. Microbes that enter the bee gut are subject to additional filters, such as low oxygen content, limited nutrient availability, and host defenses (Douglas 2015; Moran, Ochman and Hammer 2019; McLaren and Callahan 2020). Microbes within pollen provisions may have difficulty breaking open pollen grains to consume the nutrients within (Roulston and Cane 2000) or may be involved in microbe-microbe interactions. As a consequence, microbes that dominate the communities in each of these habitats may be specialized for the associated conditions (Herrera *et al*. 2010; Dhami, Hartwig and Fukami 2016; Pozo and Jacquemyn 2019; Alvarez-Perez *et al*. 2021; Christensen, Munkres and Vannette 2021), and not all microbes may be equally capable of colonizing each habitat type. Future comparative genomic or transcriptomic studies will be useful in elucidating the mechanisms involved in microbial habitat specialization (e.g., Vuong and Mcfrederick 2019).

Although its introduction was not intentional, the presence of *V*. *victoriae* provides an intriguing comparison of a microbe not adapted to this system. Despite its overwhelming presence in the fungal inoculum, *V*. *victoriae* was not able to significantly establish in any of the bee-associated habitats following its inoculation. This further supports the hypothesis that the microbes that dominate this system are specially adapted to survive in these novel environments.

Four of the focal microbes were identified across all three habitat types, which supports the hypothesis that flowers are a source of non-pathogenic microbes for bees and pollen provisions(McFrederick *et al*., 2017; Argueta-Guzmán, Spasojevic and McFrederick 2025); however, the unequal presence and abundance among habitat types also suggests environmental filtering. Larger differences in weather conditions across the two spray rounds, timing of sample collection, or differences in initial microbial communities within each hoop house could have led to the differences we saw in establishment of focal microbes across spray rounds and hoop houses. In addition, the low relative abundance of focal microbes could suggest the presence of priority effects, which have previously been found in nectar microbial communities (Toju *et al*. 2018; Chappell *et al*. 2022; Noroian, Martin, and Vannette, *unpublished data*). The samples in this study were not sterile prior to microbial inoculation, as pre-spray samples were colonized by a diverse bacterial and fungal community. The pre-existing community of microbes in each habitat type could have prevented the inoculated microbes from establishing in high numbers, potentially through direct competition for space/resources (e.g. Herrera, García and Pérez 2008; Vannette and Fukami 2016, 2018) or via modification of the environment (e.g., acidification, temperature changes, etc.; Herrera and Pozo 2010; Lee *et al*. 2019). Of the focal microbes, *Ap*. *micheneri* was found in the highest number of samples. Previous genomic analysis has revealed that *Ap*. *micheneri* is likely adapted to the bee and floral environments (Vuong and Mcfrederick 2019). For instance, *Ap*. *micheneri* contains genes which allow it to digest pollen and adhere to host guts, which could make it a better competitor (Vuong and Mcfrederick 2019). Despite these adaptations, it still did not reach high abundances in any of the sample types, further supporting the presence of priority effects.

Though the ITS86F/ITS4R primer set has been successful in previous ITS primer comparisons (Op De Beeck *et al*. 2014), the results from our fungal sequencing suggest that this primer set was biased against *M*. *reukaufii* and was unable to bind to *Au*. *pullulans* and *S*. *bombicola*. This finding also highlights the importance of including a positive control or mock community in sequencing submissions in order to maximize accuracy of results (e.g. The Human Microbiome Project Consortium 2012; Kozich *et al*. 2013; Bokulich *et al*. 2016; Karstens *et al*. 2019), as well as a reminder of the detection biases of primer sets (e.g. Bellemain *et al*. 2010; Tedersoo and Lindahl 2016). It should be noted that primer bias could have led to low abundance of our detected focal yeasts (rather than environmental filtering) and the presence of *V*. *victoriae*.

The presence of unique taxa found in both the gut and the provision indicates that flowers are not the only source of microbial acquisition for bees. Rather, adult bees may pick up microbes from other sources including their cocoons, natal nests, soil, water, and other nesting substrate (Keller, Grimmer and Steffan-Dewenter 2013; Rothman *et al*. 2019; de Sousa 2021). The environmental conditions of each habitat may contribute to the overlap in ASVs. For instance, flowers and pollen provisions contain large amounts of pollen and nectar and are both exposed to oxygen. On the other hand, adult bee guts have limited resources and typically have no or low oxygen content (McLaren and Callahan 2020). Microbes shared between bee guts, flowers, and pollen provisions would thus need to survive in these contrasting environments, which may explain the lower number of shared taxa. Instead, it seems more likely that microbes specialize within one of the habitats and use the others as temporary transport between new preferred habitats (Moran, Ochman and Hammer 2019; Vannette 2020).

When we supplemented pollen provisions with additional microbes, we found no effect on larval survival, development, pupation, or emergence. Because the pollen provisions were not initially sterile, we did not observe the survival benefits previously documented in other studies (Dharampal, Danforth and Steffan 2022). This finding has several major implications. First, it supports the use of certain microbial biocontrol agents as safe tools for managing plant pests and pathogens. For example, *Pantoaea agglomerans*, one of the microbes used in our study, has been used in the biological control of the plant pathogen fire blight (*Erwinia amylovora*) in orchard crops (e.g. (Pusey *et al*. 2011; Kim *et al*. 2012). Understanding the off-target impacts of such agents on beneficial insects is crucial to maintaining sustainable agroecosystems. Secondly, our results suggest that the native floral microbiome does not negatively impact blue orchard bee health. Current research on the interaction between bee and floral microbiome have been focused on potential implications for bee health (e.g. (Martin, Schaeffer and Fukami 2022; Brar *et al*. 2024; Nguyen and Rehan 2025). Future research should investigate how the microbiome may impact bee health under different contexts, such as in pathogen-infected or pesticide-exposed bees.

In conclusion, this study provides evidence that adult bees acquire microbes from flowers and vector them to pollen provisions for their offspring, with no detectable effect on larval health. These microbes do not need to be ingested or established in the female gut to reach the pollen provisions, suggesting that females may serve only as microbial vectors for their offspring. Flowers are often considered to be the “dirty doorknobs” of the bee world—it has long been posited and supported that pathogenic microbes are acquired from flowers, and this study finds that commensal microbes are also acquired from this transmission route. Although this study focused on a single flower species and one solitary bee species, we reveal that pollination networks serve as pathways for microbial exchange and highlights variation in microbial establishment across distinct bee-associated habitats. By directly tracking inoculated microbes through unique and complex environments, we observed the direct route of microbial transmission and differential establishment within this tractable model system. This work is broadly applicable in relation to microbial control efforts, since bees have the potential to vector insect and plant pathogens (e.g., (Durrer and Schmid-Hempel 1994; Mukhtar *et al*. 2024), but also suggests that microbial control agents could be incorporated into the pollen provision. The traditional dogma proposed by the Dutch microbiologist Martinus Wilhelm Beijerinck suggests that “everything is everywhere, but the environment selects” (O’Malley 2008); however, this study also highlights the importance of dispersal via vectors in transferring microbes between habitats. Overall, these findings suggest that factors of an environment—including pre-existing microbial communities, climate, and habitat conditions—may lead to differential transmission and establishment among microbial species.

## Acknowledgments

The authors would like to thank members of the Vannette Lab and Grace Melone for feedback on drafts of the manuscript, as well as members of the UC Davis Insect Ecology Discussion Group for feedback on the results. The authors would also like to thank Joe Tauser for assisting with cage maintenance and Dalhousie University for sequencing services.

## Data Availability

The scripts used to process this data and all data files are available on github at the following URL: https://github.com/lexienichole/Osmia-Hoophouse-2025. The sequencing data used in this study were submitted to the NCBI Sequence Reads Archive (SRA) under the Bioproject ID PRJNA1298349. All files will be made publicly available upon publication.

## Author Contributions

ANM and RLV conceived the study, inoculated flowers, and took samples. CS and NMW cared for the hoop houses and bees. ANM led lab work, data analysis, and manuscript writing. HMN contributed to lab work. RLV secured funding and contributed to manuscript writing. All authors participated in manuscript revisions.

