## Supplemental Figures for "Floral microbes provisioned by *Osmia lignaria* establish in larval food stores, but do not affect bee development or survival"

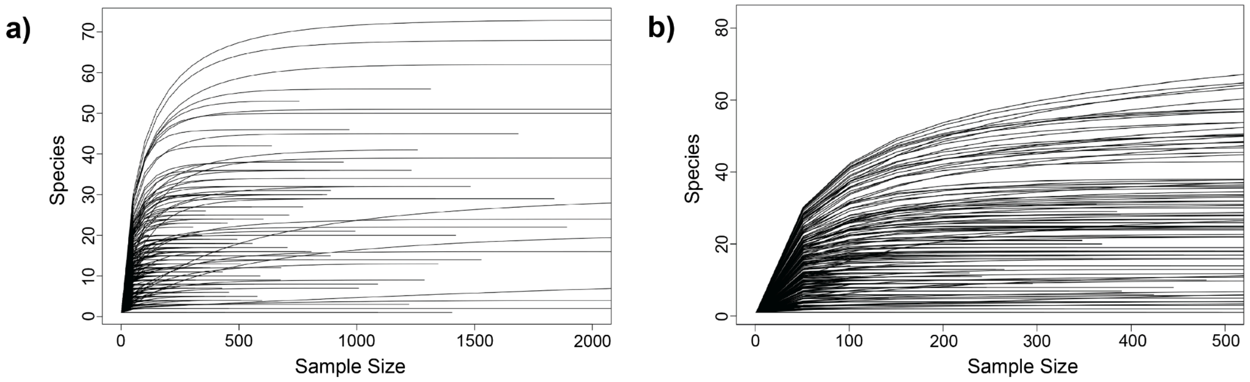


**Figure S1**: Rarefaction curves for (a) Bacteria and (b) Fungi used to determine sufficient sampling depth

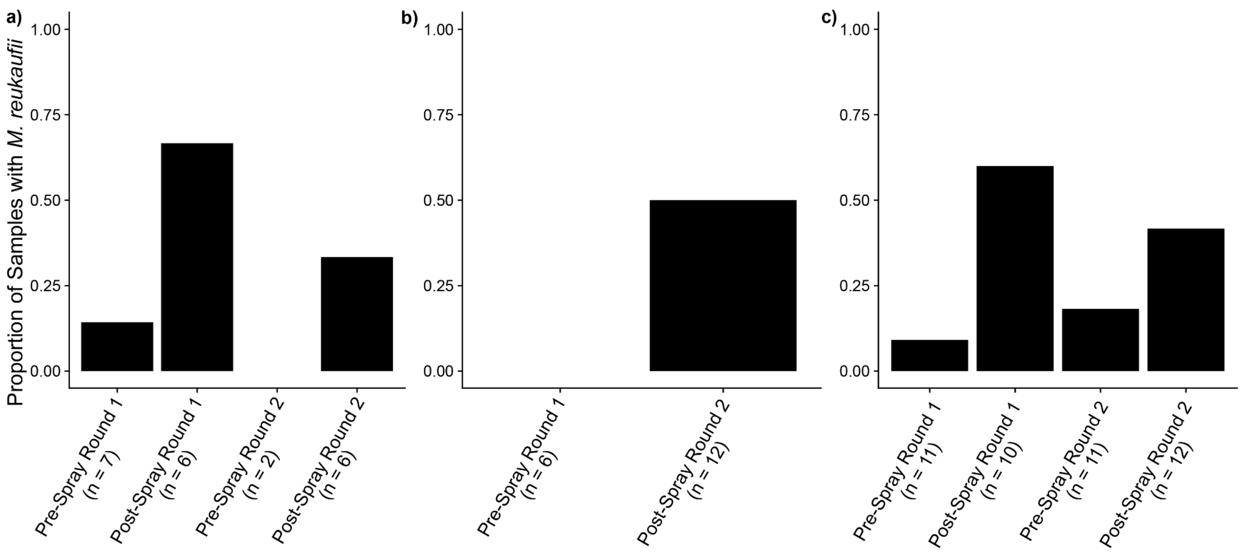


**Figure S2**: Presence of *Metschnikowia reukaufii* across (a) flowers, (b) adult bee guts, and (c) provisions detected using PCR and gel electrophoresis.

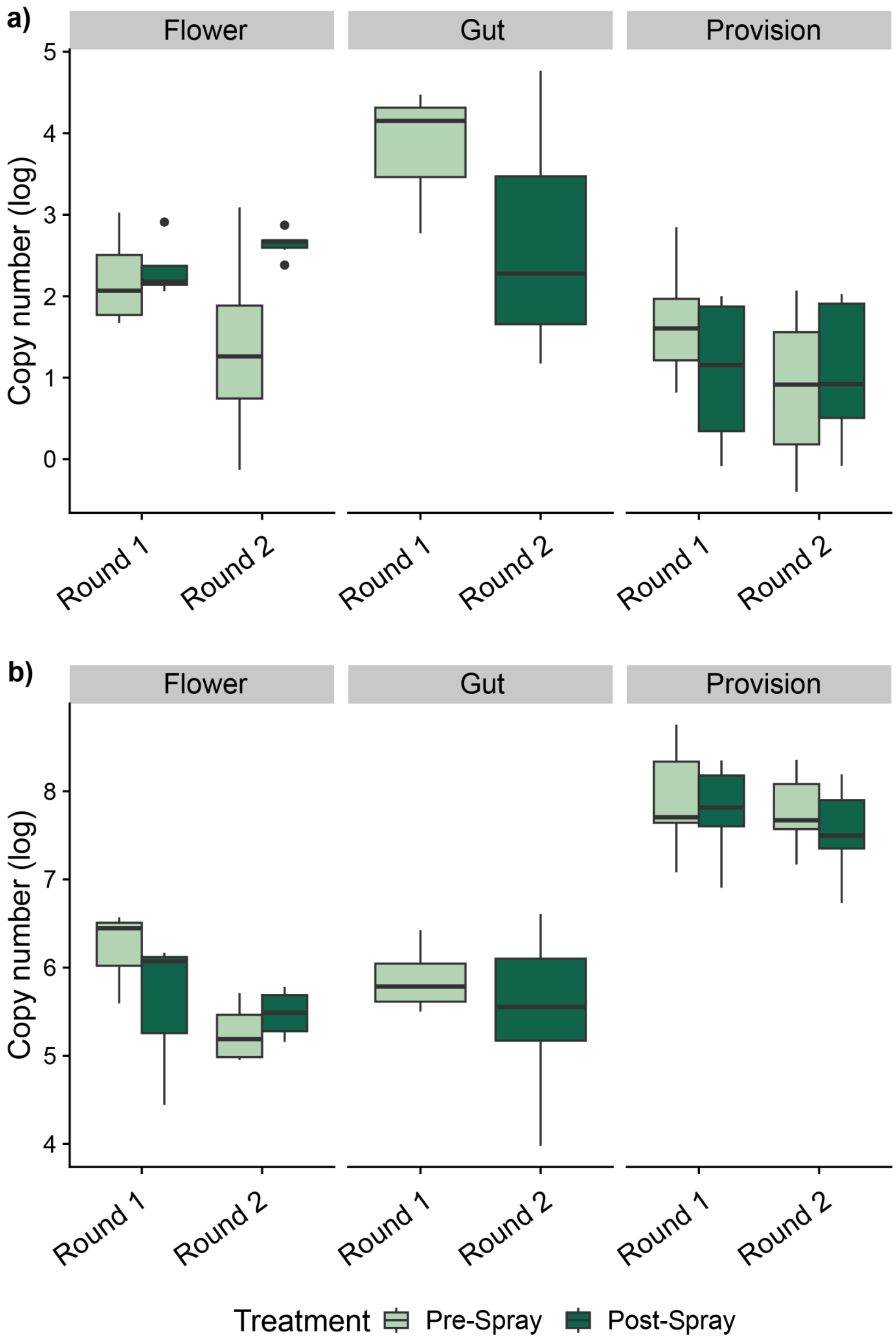


**Figure S3**: Adjusted log copy number of (a) Bacteria and (b) fungi across flowers, adult bee guts, and pollen provisions.

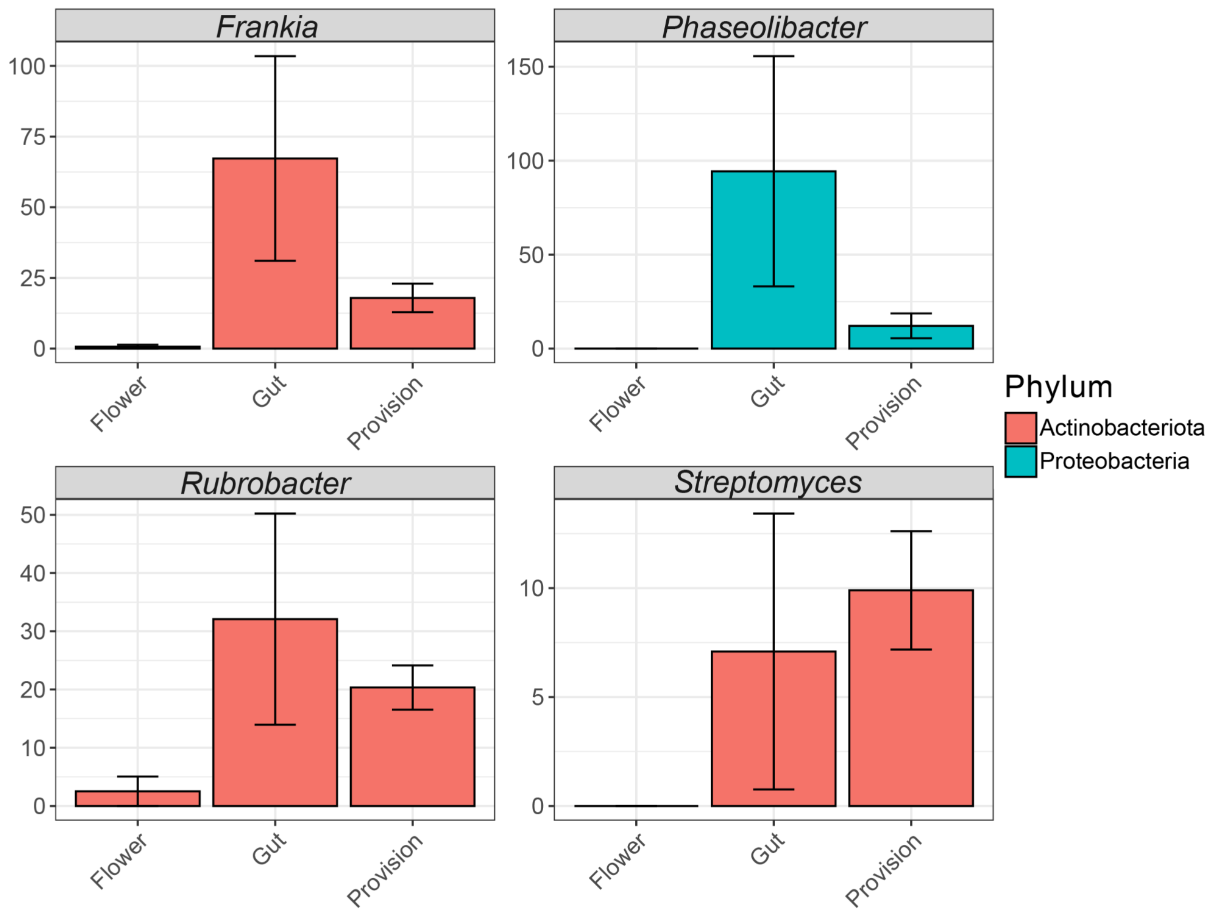


**Figure S4**: Differentially abundant bacterial genera between flowers, adult bee guts, and pollen provisions

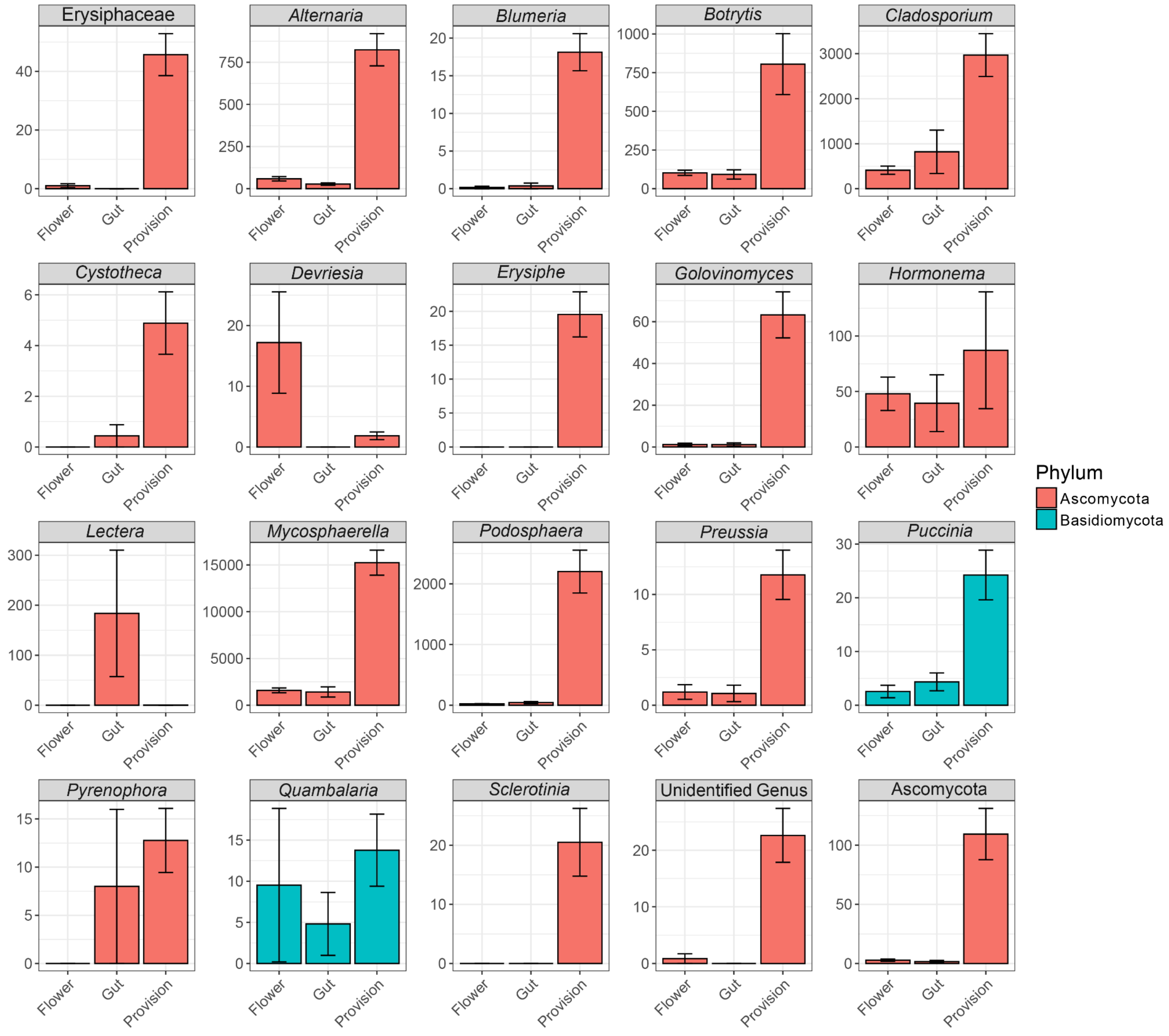


**Figure S5:** Differentially abundant fungal genera between flowers, adult bee guts, and pollen provisions
