## Supplemental Methods for "Floral microbes provisioned by *Osmia lignaria* establish in larval food stores, but do not affect bee development or survival"

### Detecting *Metschnikowia reukaufii* in samples and investigating primer bias

Because only one (*D. hansenii*) out of the four applied fungi was detected via the Illumina amplicon data, PCR and gel electrophoresis were performed on DNA extracted from all of the samples to assess potential bias in the primer set. To do this, we investigated the presence of one of the undetected focal yeasts — *M. reukaufii*. The PCR master mix was created to a reaction volume of 20 ul: 10 ul of Denville Choice Taq Blue Mastermix, 7 ul of sterile PCR water, 1 ul of the *M. reukaufii* specific forward primer Mr_LSU_F1_2 (5'-TGCAAGCAGACACAACCTCG-3'; 10uM) and 1 ul of the *M. reukaufii* specific reverse primer Mr_LSU_R1_2 (5'-CGCCAGCATCCTTGAAGAAT-3'; 10uM) (Colda *et al.*, 2021). Thermocycling conditions included initial denaturation for 10 minutes at 95 °C, 15 second denaturation at 95 °C for 40 cycles, annealing for 60 seconds at 57 °C, and elongation for 30 seconds at 72 °C (Colda *et al.*, 2021). Gel electrophoresis was performed for positive controls, negative controls, and both inocula on a 1.5% agarose gel mixed with 1% SYBR dye to ensure PCR was successful. Amplified PCR product was cleaned using EXOSAP-It enzyme following the manufacturer instructions. Samples were sent to Genewiz from Azenta Life Sciences for Sanger Sequencing, along with the inocula, two positive controls, and two negative controls. Sequence results were analyzed in SnapGene Viewer (SnapGene software (www.snapgene.com)) to confirm the presence or absence of *M. reukaufii*. If the sample failed sequencing or the sequence was dominated by Ns, it was assumed that *M. reukaufii* was not present. If the sample generated a sequence, it was searched in NCBI

BLAST to confirm it was *M. reukaufii*. To determine if *M. reukaufii* was differentially present across treatments, the same chi-squared tests as described in the Aim 1 statistical analysis methods were performed on the presence/absence dataset generated using PCR with species-specific primers.

Next, we used SnapGene to compare primer binding regions in the ITS amplified sequences. We first searched the National Center for Biotechnology Information Database for ITS sequence regions of all focal fungal taxa. Reference Sequences were as follows: NR_111252.1 for *Metschnikowia reukaufii*, NR_121483.1 for *Starmerella bombicola*, NR_144909.1 for *Aureobasidium pullulans*, NR_073260.1 for *Vishnicozyma victoriae*, and NR_120016.1 for *Debaryomyces hansenii*. For each sequence, we first attempted to add the forward and reverse primers to investigate if the primers would bind. Then, we aligned the forward and reverse primer sequences with the microbial sequence. We noted whether SnapGene found a primer binding site, and the number of different bases between the primers and the microbial sequence.
